# Baculovirus utilizes cholesterol transporter Niemann–Pick C1 for host cell entry

**DOI:** 10.1101/312744

**Authors:** Zhihong Li, Junhong Wei, Youpeng Fan, Xionge Mei, Qiang He, Yonghua Zhang, Tian Li, Mengxian Long, Jie Chen, TongBao Liu, Jialing Bao, Zhonghuai Xiang, Guoqing Pan, Chunfeng Li, Zeyang Zhou

## Abstract

The dual roles of baculovirus for the control of natural insect populations as an insecticide, and for foreign gene expression and delivery, have called for a comprehensive understanding of the molecular mechanisms governing viral infection. Here, we demonstrate that the *Bombyx mori* Niemann-Pick C1 (BmNPC1) is essential for baculovirus infection in insect cells. Both pretreatment of *Bombyx mori* embryonic cells (BmE) with NPC1 antagonists (imipramine or U18666A) and down-regulation of NPC1 expression resulted in a significant reduction in baculovirus BmNPV (*Bombyx mori* nuclear polyhedrosis virus) infectivity. Furthermore, we show that the major glycoprotein gp64 of BmNPV, responsible for both receptor binding and fusion, is able to interact predominantly with the BmNPC1 C domain, with an enhanced binding capacity at low pH conditions, indicating that NPC1 most likely plays a role during viral fusion in endosomal compartments. Our results, combined with previous studies identifying an essential role of hNPC1 in filovirus infection, suggest that the glycoprotein of several enveloped viruses possess a shared strategy of exploiting host NPC1 proteins during virus intracellular entry events.

**IMPORTANCE:** BmNPV is one of the most important members of the *Baculoviridae*; many viruses in this family have been frequently employed as viral vectors for foreign gene delivery or expression and as biopesticides, but their host receptors still remain unclear. Here, we describe that the intracellular cholesterol transporter BmNPC1 is indispensable for BmNPV infection in insect cells, and it interacts with the major viral glycoprotein gp64. Our study on the role of BmNPC1 in baculovirus infection has further expanded the list of the enveloped viruses that require host NPC1 proteins for entry, and will ultimately help us to uncover the molecular mechanism of the involvement of NPC1 proteins in the entry process of many enveloped viruses.

---

Enveloped viruses include a number of pathogens with significant importance to animal or human health. As a typical double-stranded DNA enveloped virus, baculoviruses are known to infect invertebrates, with over 600 host species described (1, 2). Due to a strong species-specific tropism for arthropods, baculovirus has been widely used as a biopesticide (3). Concurrently, baculovirus can transduce a broad range of vertebrate cells, including human, bovine, fish avian, and even primitive cells such as embryonic stem cells, warranting its application as a biological tool for gene delivery (4–6). During its viral infection cycle, baculovirus produces two virion phenotypes: occlusion-derived virus (ODV) which transmits to insects by the oral route and specializes to infect midgut epithelial cells, and budding virus (BV) produced during nucleocapsid budding from the basolateral membrane of infected cells, responsible for systemic host infection (7). During viral infection, BV first attaches to the host cell surface, and is internalized by endocytosis to late endosomes (8); subsequently, the viral membrane of BV fuses with the endosomal membrane triggered by low pH and nucleocapsids are released into cytoplasm followed by viral replication (9, 10). The major BV envelope protein, glycoprotein 64 (gp64), has been shown to be essential for both virus attachment and membrane fusion (11). Despite the fact that a number of host molecules including heparin sulfate (12), phospholipids (13) or *Bombyx mori* receptor expression-enhancing protein (BmREEPa) (14) have been identified to be involved in BV attachment or binding, the exact identity of host receptors for baculovirus still remains elusive.

Among numerous baculoviruses, *Bombyx mori* nuclear polyhedrosis virus (BmNPV) is one of the most frequently employed in the research. It has been reported previously that BmNPV infection was enhanced by *Bombyx mori* promoting protein (BmPP), a family member of Niemann-Pick C2 (NPC2), in silkworm BoMo cells (15). The addition of BmPP to culture media at a concentration of 1 μg /ml resulted in a 1000–10,000-fold increase of BmNPV production; however, the detailed molecular mechanism has not yet been fully elucidated. Human NPC2 protein functions as a central shuttle binding to Niemann-Pick C1 (NPC1) to export low density lipoprotein (LDL)-derived cholesterol from late endosomes and lysosomes to other cellular compartments (16). The role of BmPP in promoting BmNPV infection indicates that host proteins responsible for cholesterol transport may be involved in baculovirus infection. Interestingly, the infection of Ebola virus (EBOV) has proven to be dependent on cholesterol transporter NPC1, which serves as a virus intracellular receptor for filovirus entry (17–19). NPC1 is a ubiquitously expressed membrane protein which is mainly involved in intracellular cholesterol transport (20). Structurally, NPC1 is comprised of three large luminal ‘loop’ domains, A, C, and I (21, 22). Domain A, also called the N-terminal domain (NTD), is involved in cholesterol binding and exporting cholesterol from lysosomes into the cytosol. Domain C directly and specifically binds to NPC2 and constitutes the scaffold to properly accommodate NPC2 for hydrophobic handoff of cholesterol to the pocket of NTD (16, 23, 24). Furthermore, NPC1 domain C is able to bind to the cathepsin-primed form of Ebola glycoprotein (GPcl) (23, 25); the absence of domain C results in complete resistance to infection by EBOV, indicating that this domain is essential for EBOV entry. The third luminal domain I interacts with domain C, and may play a supporting role in cholesterol transport (23).

The recent findings of NPC1 protein as an intracellular receptor for filovirus, coupled with prior evidence supporting the involvement of NPC2 (BmPP) in BmNPV infection, prompted us to investigate the potential role of NPC1 homologs in BmNPV infection in insect cells. Here, we demonstrate that BV of BmNPV infection requires the expression of BmNPC1. BmE cells pre-treated with the NPC1 inhibitors imipramine or U18666A, which can mimic the molecular phenotype of NPC disease (26, 27), results in marked inhibition of viral replication and production. Silencing, knocking out of BmNPC1 expression or blocking BmNPC1 function with domain-specific antibodies also substantially impairs BmNPV proliferation *in vitro*. Together, these data show that BmNPC1 is an essential host factor for baculovirus infection in insect cells, and identifies a shared strategy of exploiting the host NPC1 protein by a group of enveloped viruses during virus intracellular entry events.

## RESULTS

### Identification of *npc*1 homologs in the *Bombyx mori* genome

To identify *Bombyx mori* homologs of the *npc1* gene, the human NPC1 (hNPC1) gene sequence (gi|255652944) was used to conduct a BLAST search in the *Bombyx mori* genome database in NCBI. Results revealed that the gene (XP_012544312.1) encoding a protein of 1334 amino acids long shared the highest sequence similarity of 43% with hNPC1 (Fig S1), indicating this gene is a potential homolog of a vertebrate NPC1 family member. This gene was referred as BmNPC1 in our study. Similar to hNPC1, structure prediction revealed that BmNPC1 protein contains 13 transmembrane-spanning helices and 3 large luminal loops, namingly domain A (residues 1 to 270), domain C (residues 408 to 645), and domain I (residues 881 to 1136) (Fig 1A); Like hNPC1, BmNPC1 protein includes 3 domains: the NTD domain (NPC1 N-terminal domain, residues 31 to 270), the SSD domain (Sterol-sensing domain, residues 674 to 828), and the patched domain (residues 1088 to 1289). Phylogenetically, BmNPC1 can be grouped in a single clade with *Apis cerana*, *Drosophila melanogaster*, and is relatively distant from NPC1s of *Saccharomyces cerevisiae* and *Toxoplasma gondii* (Fig S1).

**FIG 1.**
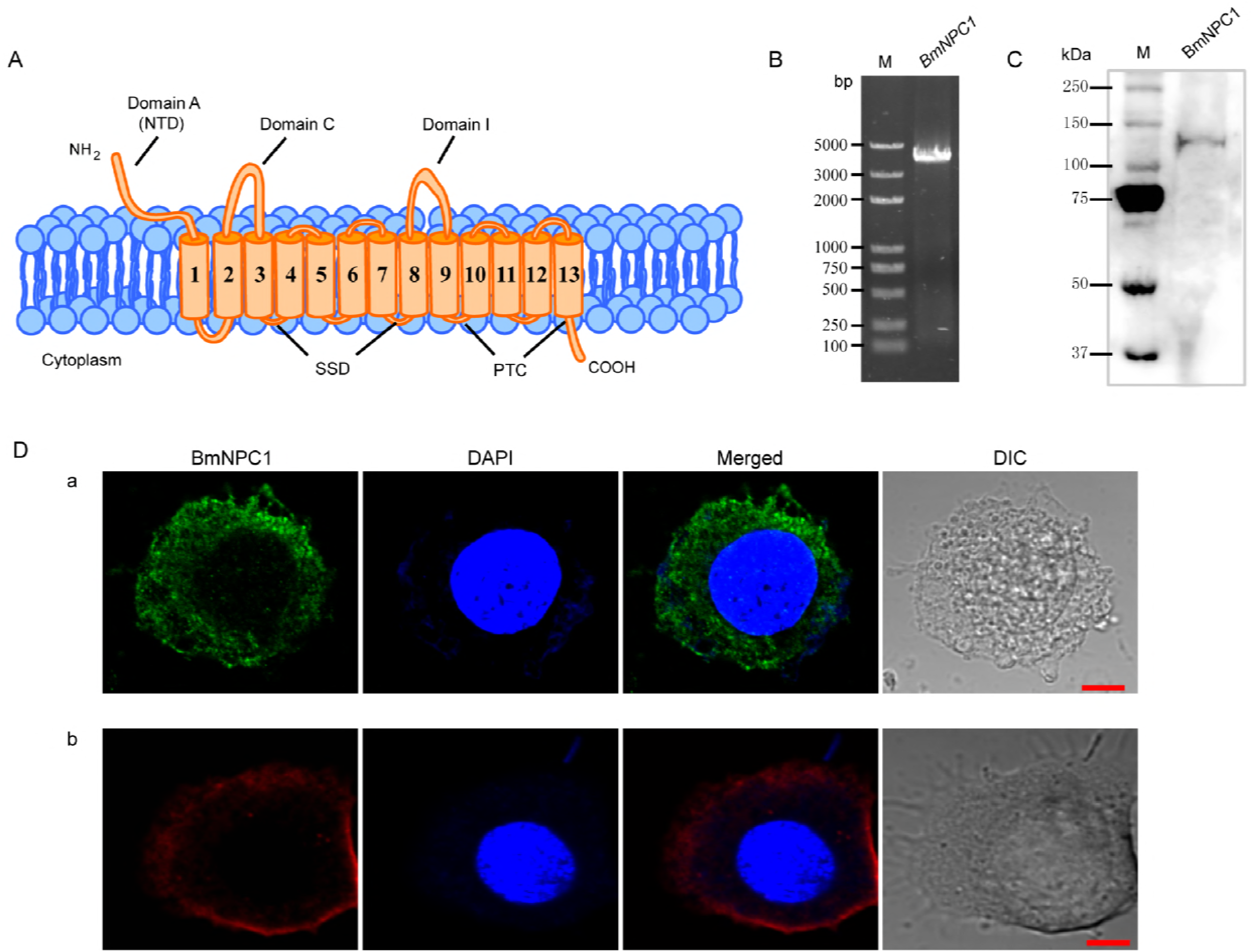
Expression and localization of BmNPC1 in BmE cell. (A) Schemes show the main structural characteristics of BmNPC1. BmNPC1 contained 13 transmembrane helices and 3 large conservation domains: N-terminal domain (NTD), sterol sensing domain (SSD) and patched domain (PTC). The three large luminal loops were named as domains A, C, and I. (B) BmNPC1 full-length gene was PCR amplified with specific primers (supplementary Table S1) from BmE cell cDNA. (C) Immunoblot of BmNPC1 in BmE cells with PcAb-hNPC1 antibody. (D) Localization of BmNPC1 in BmE cells. BmE cells were permeabilized (a) and unpermeabilized (b) followed by immunostaining with PcAb-hNPC1 antibody. The vesicle lumen and membranes were labeled by PcAb-hNPC1 antibody in the permeabilized BmE cells (a1, green fluorescence). The plasma membrane was labeled by PcAb-hNPC1 antibody in the unpermeabilized BmE cells (b1, red fluorescence). The nucleus was stained by DAPI. Bar, 5 μm.

To further study the biological function of BmNPC1 and its role in baculovirus infection in insect cells, the full length (4 kb) coding sequence (CDS) of BmNPC1 was cloned from the *Bombyx mori* cDNA library (Fig 1B); the sequence was 100% identical to the sequence of XP_012544312.1 published in the NCBI database. Furthermore, a rabbit polyclonal antibody against hNPC1 (PcAb-hNPC1) was used to determine protein expression and localization of BmNPC1 in BmE cells. A single band of ~140 kDa was detected in BmE whole cell lysate by Western blot (Fig 1C). BmNPC1 proteins were localized in the plasma membrane, vesicle lumen and membranes in BmE cells as revealed by IFA (Fig 1D). Taken together, we demonstrate that the putative BmNPC1 gene (XP_012544312.1) shares high sequence and structural similarity with hNPC1 and was transcribed and translated in BmE cells, with the protein abundantly expressed in the plasma membrane and intracellular vesicles.

### The potent NPC1 antagonists, imipramine and U18666A, inhibited BmNPV infection in BmE cells

To investigate whether BmNPC1 is involved in BmNPV infection in insect cells, we first examined the effect of two small molecule NPC1 antagonists on BmNPV-GFP infection in BmE cells. Both small molecules (imipramine and U18666A) have previously been shown to mimic the molecular phenotype of NPC disease and block the exit of cholesterol from late endosomal compartments (17, 27). BmE cells were pretreated with imipramine (25-100 μM for 2 h) or U18666A (0.5 to 10 μM for 24 h) before the addition of BmNPV viral inoculum. The effect of drug pretreatment on viral infection was evaluated based both on the percentage of GFP expression positive cells at 72 h post infection (p.i.), and the quantification of viral DNA accumulation in infected cells by qPCR. Imipramine and U18666A showed no evident cytotoxicity in BmE cells (Fig S2), and drug-pretreatment resulted in a dose-dependent reduction in viral infectivity as demonstrated by the percentage of GFP positive cells (Fig 2A and 2B). The reduction in GFP expressing positive cells upon drug treatment was statistically significant at concentrations of 25-100 μM for imipramine and of 1-10 μM for U18666A (Fig 2C, 2E). Consistent with the quantification of GFP positive cells, the viral load (represented by relative copy number of gp41) decreased significantly at concentrations of 50-100 μM for imipramine and at concentrations of 1-10 μM for U18666A. For BmE cells pretreated with 100 μM of imipramine or 10 μM of U18666A, the viral load was reduced to 27% and 10%, respectively, compared to the vehicle along treatment control (Fig 2D, 2F). In summary, we conclude that blockage of BmNPC1 function by small molecules can efficiently reduce BmNPV infection in BmE cells, indicating that functional BmNPC1 is required for BmNPV infection in insect cells.

**FIG 2.**
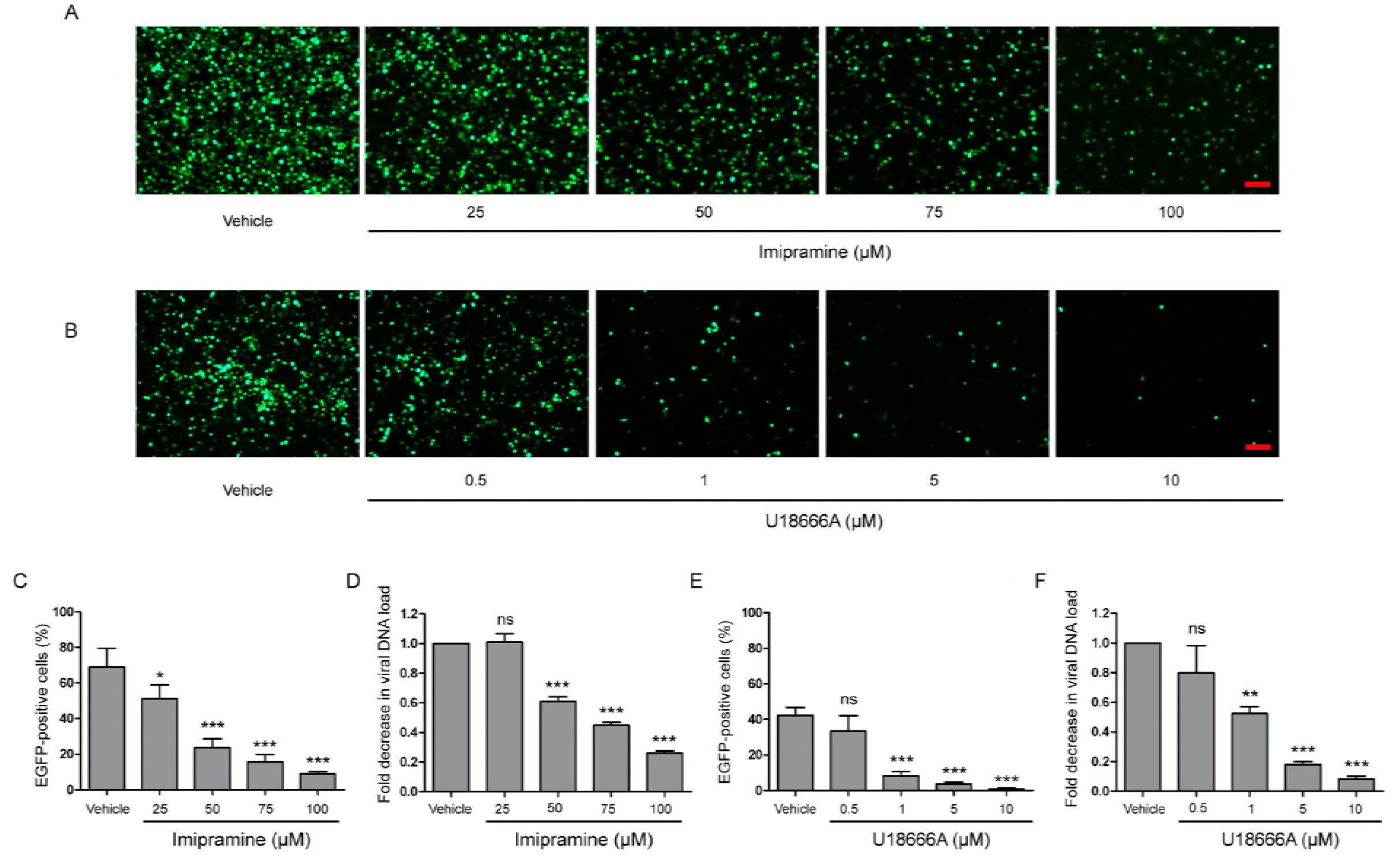
Cationic amphiphilic compounds imipramine or U18666A inhibits BmNPV replication in BmE cell. BmE cells were pretreated with imipramine (A) or U18666A (B) for 2 and 24 h, respectively followed by GFP-BmNPV infection at an MOI of 1 for 72 h. Viral infection was indicated by positive GFP expression in BmE cells. Bar, 200 μM. Quantification of GFP-BmNPV infectivity in BmE cells pretreated with impiramine (C) or U18666A (E). The percentage of GFP positive cells was counted from at least five different fields. Fold decrease of viral DNA load in BmE cells pre-treated with imipramine (D) or U18666A (F) compared to control. The viral DNA load was analyzed by qPCR analysis using gp41 DNA primers, and the viral DNA load in cells treated with blank solvent was set as “1”. Relative copy numbers were calculated using *B. mori* GADPH DNA as the internal control. Independent-samples T tests were used for statistical analysis. For panels C-F, \**P*<0.05; \*\**P*<0.01; \*\*\**P*<0.001; ns, non-significant, Means ± s.d. are shown (n=3).

### Knockdown of BmNPC1 expression by RNAi reduced virus infection in BmE cells

To further study the role of BmNPC1 in BmNPV infection, two sets of RNAi vectors targeting BmNPC1 (psl-BmNPC1-a and psl-BmNPC1-b) were constructed (Fig 3A), and independently transfected into BmE cells. The reduction of BmNPC1 expression at the mRNA level following BmNPC1-RNAi treatment was detected by qPCR at various time points post transfection (p.t.). psl-BmNPC1-b was selected to knock down BmNPC1 expression in BmE cells due to greater efficiency and duration in reducing the expression of BmNPC1 (40% of reduction compared to psl-null at 48 h, with the knock-down effect persisting until 96 h p.t.) (Fig 3B). BmE cells were treated with psl-BmNPC1-b or psl-null for 48 h prior to infection with BmNPV-GFP. As shown in Fig 3C and 3D, viral infectivity as indicated by GFP expression was decreased significantly in psl-BmNPC1-b treated cells at 48, 72 and 96 h p.i. compared to that in control RNAi treated cells. Furthermore, viral loads represented by relative copy number of gp41 in BmNPC1 RNAi treated cells was less than 10% of that in control RNAi-treated cells at 72 h p.i. (Fig 3E). The viral titer (5.6 log_10_TCID_50_/ml) at 96 h p.i. as quantified by TCID_50_ in BmNPC1 RNAi treated cells was significantly lower compared to that in control cells (10.8 log_10_TCID_50_/ml, Fig 3F). Collectively, we conclude that knocking down BmNPC1 by RNAi expression in BmE cells can substantially reduce BmNPV infectivity.

**FIG 3.**
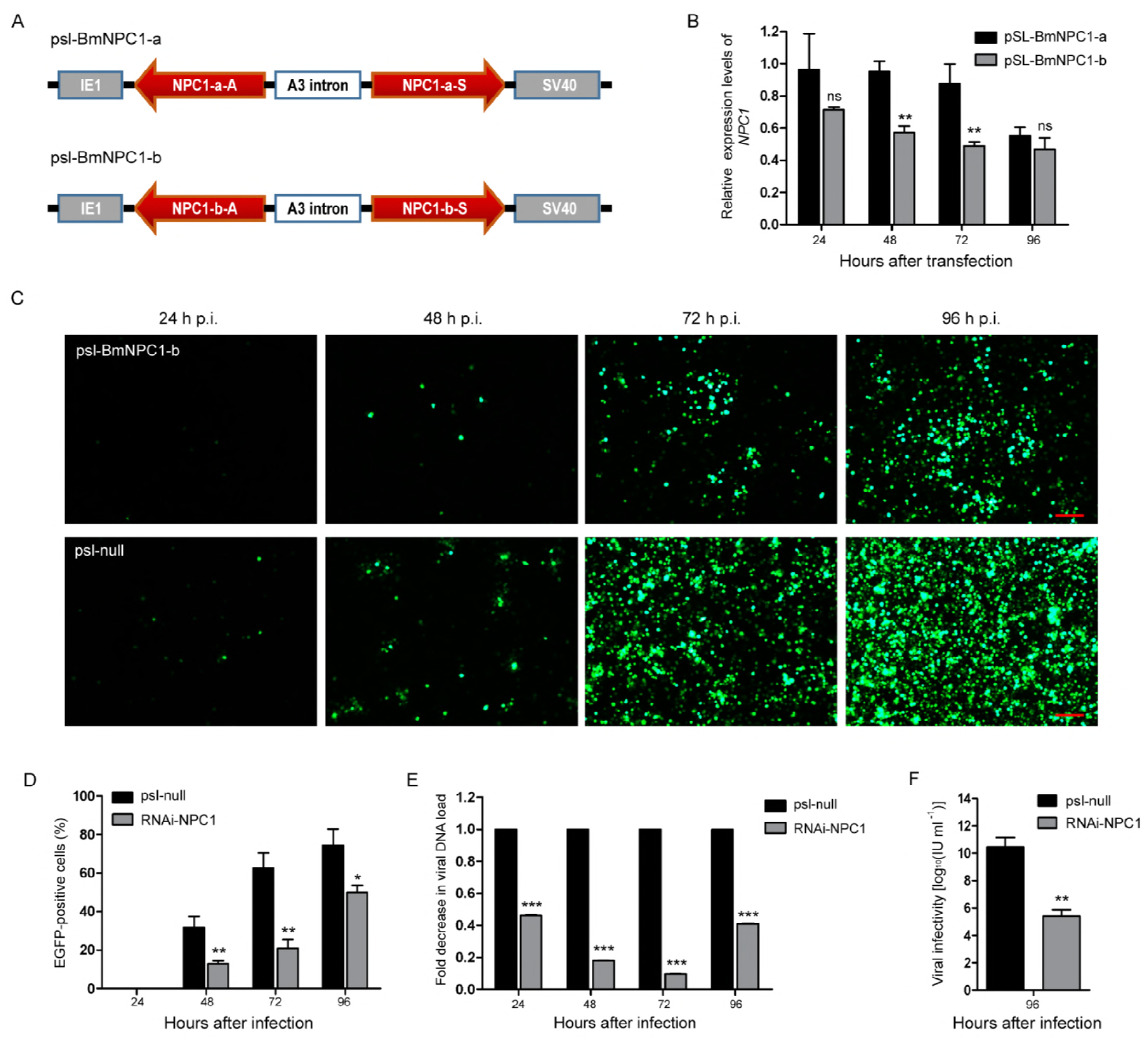
Silence of BmNPC1 inhibits BmNPV replication in BmE cell. (A) Schematic diagram of the BmNPC1 RNAi vector construction. IE1 indicates the *ie1* promoter of BmNPV. NPC1-a and NPC1-b are two NPC1 fragments, NPC1-a-S, NPC1-b-S, NPC1-a-A and NPC1-b-A represents the sense fragment (S) and the antisense fragment (A) of NPC1-a and NPC1-b, respectively. These backbones were inserted into pSL1180 vector (‘‘head to head’’), the arrow direction of gene fragment represented 5’– 3’. The A3 intron was a spacer cloned from the genomic DNA of *B. mori Dazao*. SV40 was the polyadenylation signal. (B) The effect of BmNPC1 RNAi transfection in BmE cell. BmE cell was transfected with psl-BmNPC1-a or psl-BmNPC1-b, and the relative expression levels of *NPC1* was detected by qPCR at the indicated times. (C) Fluorescence microscopy of BmNPV infected BmE cells transfected with psl-BmNPC1-b or null RNAi at indicated times. Cells transfected with psl-BmNPC1-b or null RNAi were infected with BmNPV at an MOI of 3. BmNPV infection positive cells were indicated by GFP expression. Bar, 200 μM. (D) Quantification of BmNPV infection based on GFP expression in BmE cells transfected with psl-BmNPC1-b and psl-null RNAi vector. The percentages of GFP positive cells of five different fluorescent fields were counted in each group. (E) The fold change of viral DNA load in cells transfected with psl-BmNPC1-b and psl-null RNAi vector. The viral DNA load were analyzed by qPCR analysis using gp41 DNA primers, the viral DNA loads in cells transfected with psl-null RNAi vector at the indicated times were set as “100%”. Relative copy numbers were calculated using *B. mori* GADPH DNA as the internal control. (F) Virus titer determination by TCID_50_ end point dilution assays. Supernatants were harvested at 96 h p.i. from the cells infected with BmNPV for the virus titer titration. Virus infection was determined by positive GFP expression in BmE cells by fluorescence microscopy. For panels B, D-F, \**P*<0.05; \*\**P*<0.01; \*\*\**P*<0.001; ns, non-significant, Means ± s.d. are shown (n=3).

### BmNPC1 null stable cell line was resistant to BmNPV infection

To further confirm that BmNPC1 is essential for BmNPV infection, we generated a BmNPC1 null stable cell line by using CRISPR-Cas9 technology. Partial BmNPC1 gene was replaced by *ie1*-DsRed and A3-*neo* expression cassettes (Fig S3B), and the mutated BmNPC1 gene was confirmed by PCR amplification and sequencing (Fig S3D). We next tested the susceptibility of the NPC1-mutant cells to BmNPV-GFP virus infection, and found that the percentage of GFP positive cells among BmNPC1-null cells as indicated by the expression of red fluorescence was less than 2% of that of the control cells at 96 h p.i. (Fig 4A and 4B). Concurrently, the virus load as evaluated by relative copy number of gp41 was reduced to 20% of that in control cells (Fig 4C). Altogether, these data confirmed that BmNPC1 is required for BmNPV infection, and the cells lacking functional BmNPC1 exhibit substantially reduced susceptibility for BmNPV infection.

**FIG 4.**
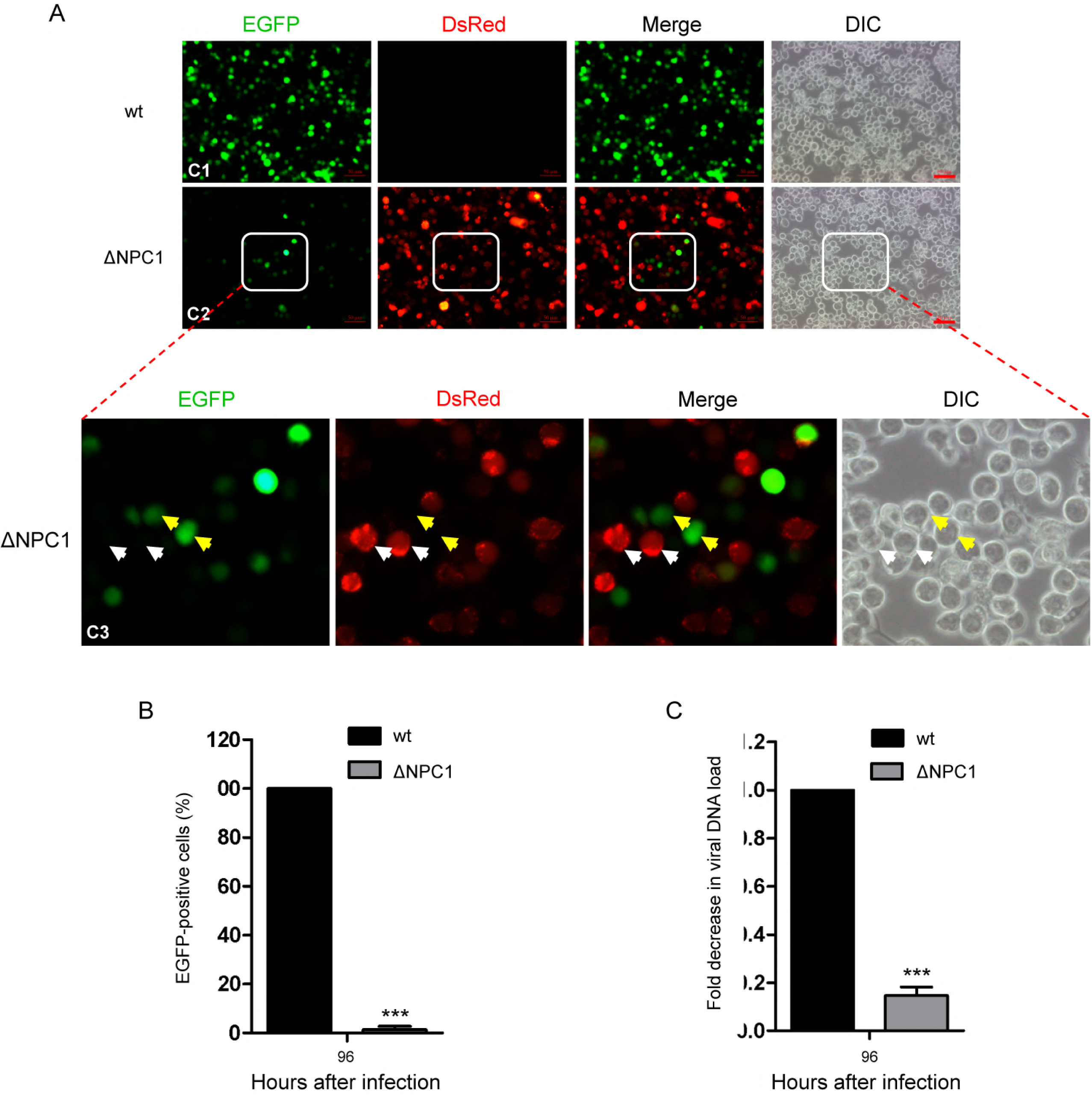
Knock out BmNPC1 by CRISPR-Cas9 system inhibits BmNPV replication in BmE cell. (A) Fluorescence microscopy of BmNPV infected wild-type and BmNPC1 knocked out BmE cells. BmNPV infection positive cells were indicated by GFP expression and BmNPC1 knocked out BmE cells were indicated by DsRed expression. Bar, 100 μM. (B) Quantification of BmNPV infection based on GFP expression in BmE cells. The percentages of GFP positive cells of five different fluorescent fields were counted in each group and the means were shown in bar graph. Three independent experiments were carried out. (C) Quantification of BmNPV infection based on viral DNA load in BmE cells. The viral DNA load were analyzed by q-PCR analysis using gp41 DNA primers, the viral DNA loads in wild-type cells at the indicated times were set as “100%”. Relative copy numbers were calculated using *B. mori* GADPH DNA as the internal control. For panels B, C, \*\*\**P*<0.001; ns, non-significant, Means ± s.d. are shown (n=3).

### BmNPV gp64 protein interacted with BmNPC1 mainly via C domain

It has been shown previously that hNPC1 is able to interact with a primed form of Ebola virus glycoprotein GP, the primary glycoprotein responsible for receptor binding and fusion (18, 25). To investigate whether BmNPC1 contributes to BmNPV infection by interacting with glycoprotein gp64 during viral entry, we performed Co-immunoprecipitation with in-vitro expressed proteins (BmNPC1 domain A, C, I and BmNPV gp64). Gp64-attached protein G agarose beads through mouse anti-gp64 antibody were incubated with BmNPC1 domain-specific proteins (domain A, C and I, respectively) for immunoprecipitation. The proteins that bound to the beads, i.e. the co-immunoprecipitation samples were then analyzed by Western blot. As shown in Fig 5A, gp64 proteins could be detected in the immune pellets by gp64 monoclonal antibody. The same immune pellets were subsequently probed with anti-HA antibody, and a specific band corresponding to the NPC1 domain A, C and I in each individual blot was shown in Fig 5B, and the band corresponding to NPC1-C displaying the strongest signal. Our results suggest that gp64 is able to interact with distinct domains (A, C and I) of the NPC1 protein, with the strongest binding affinity for NPC1-C. Next, we performed the reciprocal co-immunoprecipitation experiments, in which NPC1 domain specific protein-attached beads were used to immune-precipitate gp64 proteins. The presence of NPC1-A, C or I in the retrieved beads was confirmed with anti-HA antibody (Fig 5C), however, the band corresponding to gp64 was only present in the NPC1-C-attached immune pellets (Fig 5D). Our results suggest that domain C of BmNPC1 is sufficient for gp64 binding, while in comparison, the binding affinity of BmNPC1-A or BmNPC1-I to gp64 is relatively low.

**FIG 5.**
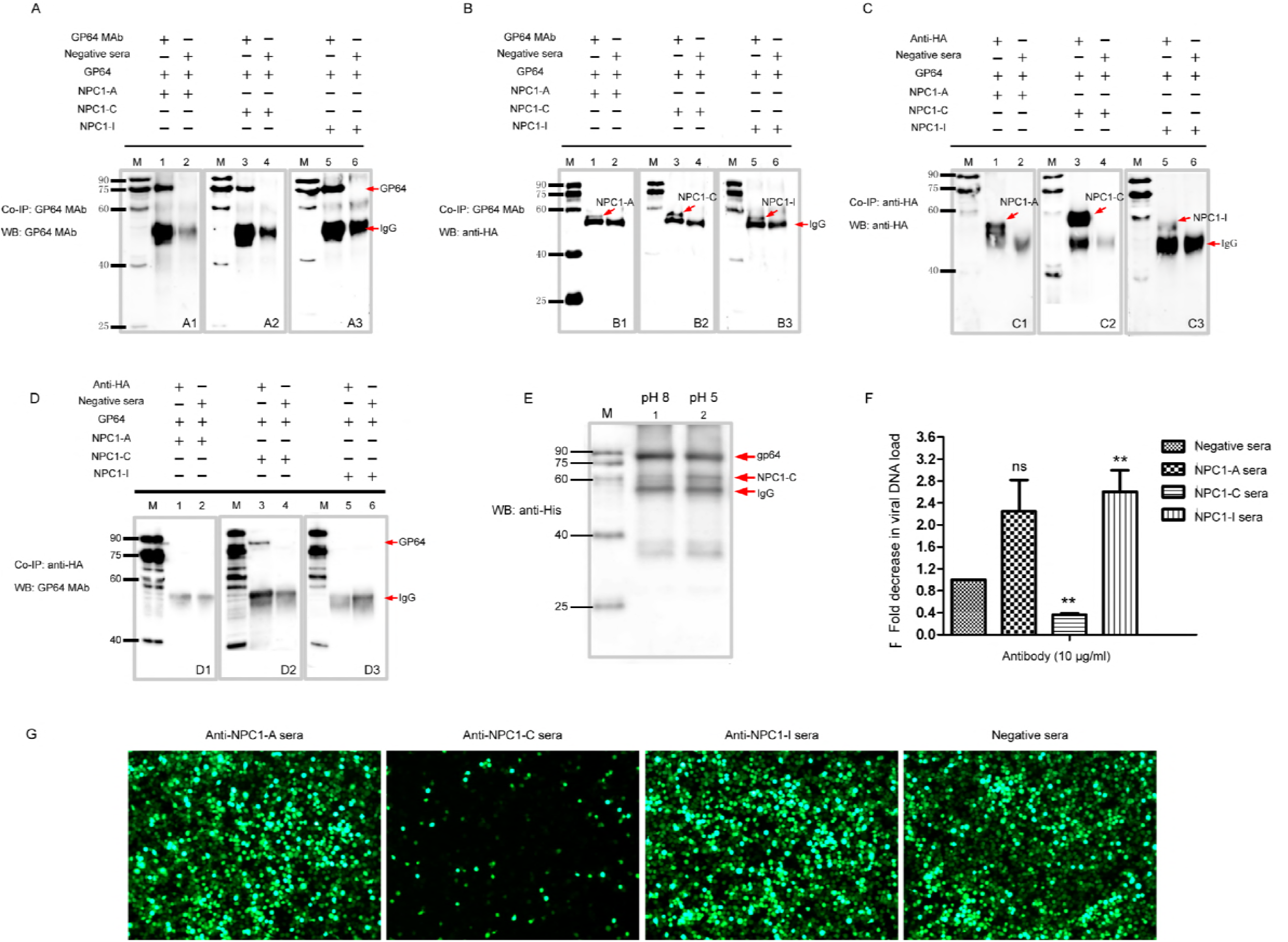
BmNPC1-C interacts with BmNPV gp64 and the antiBmNPC1-C antibody pretreatment reduces BmNPV infection. (A, C) Western blotting of co-immunoprecipitate samples was used to analyze the interaction between BmNPV gp64 and BmNPC1 domain-specific proteins. BmNPC1 domain A, C, I and gp64 were expressed by In Vitro Translation Kit (see Materials and Methods). Then the BmNPC1 domain-specific proteins were incubated with gp64, respectively, and coimmunoprecipitated with anti-gp64 monoclonal antibody. For the three immune pellets, anti-gp64 monoclonal antibody was used to detect gp64 protein (A). A1, gp64 incubated with BmNPC1-A; A2, gp64 incubated with BmNPC1-C; A3, gp64 incubated with BmNPC1-I. Anti-HA antibody was used to detect BmNPC1-A, BmNPC1-C, or BmNPC1-I at the corresponding immune pellets (C). C1, gp64 incubated with BmNPC1-A; C2, gp64 incubated with BmNPC1-C; C3, gp64 incubated with BmNPC1-I. Lane M, EasySee Western marker. (B, D) BmNPC1 interacted with BmNPV gp64 assay by CoIP. Anti-HA monoclonal antibody was used for IP with BmNPC1-A, BmNPC1-C, or BmNPC1-I (B). B1, BmNPC1-A incubated with gp64; B2, BmNPC1-C incubated with gp64; B3, BmNPC1-I incubated with gp64. The anti-gp64 antibody was used for CoIP with gp64 (D). D1, BmNPC1-A incubated with gp64; D2, BmNPC1-C incubated with gp64; D3, BmNPC1-I incubated with gp64. Lane M, *EasySee* Western marker. (E) The pH dependent interaction between gp64 and BmNPC1-C. The mixtures of NPC1-C and gp64 were coimmunoprecipitated with anti-gp64 antibody, and anti-His antibody was used for immunoblot. Red arrows indicated the corresponding proteins showed at the arrows back. Lane M, EasySee Western marker. Line 1, pH 8; Line 2, pH 5. (F) The fold change of viral DNA load in BmE cells preincubated with domain specific BmNPC1 antibodies. BmE cells preincubated with anti-NPCI-A, C, I domain specific antibody or control serum at a concentration of 10 μg/ml for 2 h before cells were infected with BmNPV at an MOI of 3 for 2 h, and the viral DNA loads were analyzed by qPCR using gp41 DNA primers. The viral DNA load in cells preincubated with negative serum was set as “1”. Relative copy numbers were calculated using *B. mori* GADPH DNA as the internal control. \*\**P*<0.01; ns, non-significant, Means ± s.d. are shown (n=3). (G) Fluorescent microscopy of BmNPV infected BmE cells preincubated with domain specific BmNPC1 antibodies. BmE cells were treated and infected as described in (F). The cells were imaged by fluorescence microscopy at 72 h p.i. The positive GFP expression indicates BmNPV infected cells.

To further confirm these interactions, we employed a yeast two-hybrid system (Y2H). Full-length gp64 was cloned into the pGBKT7 vector as bait, and BmNPC1-A, BmNPC1-C, BmNPC1-I were used as the prey. These Y2H screens revealed that BmNPC1-C interacted with full-length gp64, whereas BmNPC1-A or BmNPC1-I bound to gp64 with a weak affinity (Fig S4). Collectively, these results demonstrate a specific interaction between BmNPC1 domain C and BmNPV gp64.

Next, we investigated whether the interaction between BmNPV gp64 and BmNPC1 domain C was pH dependent. For this purpose, the Co-IP between BmNPV gp64 and BmNPC1-C was performed at pH 5 or 8. NPC1-C and gp64 proteins were expressed with His tag in vitro, and then the protein mixtures were incubated with protein G beads that had been conjugated with mouse monoclonal antibody to gp64. Anti-His antibody was used to detect both gp64 and NPC1-C at the same blot for more accurate comparison. As shown in Fig 5E, we found a stronger BmNPC1-C signal at pH 5 compared to that at pH 8, indicating that BmNPV gp64 binding affinity to BmNPC1-C was enhanced at a lower pH environment.

To further confirm which domain of BmNPC1 was most critical for BmNPV infection, we assessed the susceptibility of BmNPV infection in the presence of BmNPC1 domain specific antibodies. Specific antibodies targeting extracellular loops A, C and I of BmNPC1 were produced and incubated with BmE cells for 2 h prior to virus infection. Compared to cells treated with a negative IgG control antibody, BmNPV infection was significantly reduced in BmE cells treated with antibodies specific for BmNPC1-C, but not BmNPC1-A or BmNPC1-I as measured by GFP positive cells and viral gene expression by qPCR (Fig 5F and 5G). These data suggest that the interaction between BmNPC1 domain C and BmNPV gp64 is essential for BmNPV infection in BmE cells.

## DISCUSSION

Baculoviruses have been widely used as important biological agents for controlling insect populations, and powerful biological tools for gene delivery and expression; a better understanding of the molecular mechanisms and host factors involved in baculovirus virus entry is of great significance in bioscience and biotechnology. In this study, we used the well-studied baculovirus BmNPV as a tool to investigate the host factor requirements for baculovirus infection, and found that BmNPC1, the hNPC1 homolog in insect cells, is indispensable for BmNPV infection in insect cells. Our results, together with previous work identifying a role for hNPC1 as an intracellular receptor for Ebola virus entry, have revealed that the conserved host protein NPC1, essential for cholesterol homeostasis, has been exploited by a group of divergent viruses for entry.

As an essential component of cellular membranes, cholesterol is one of the most important lipids for maintaining cell viability, cell signaling and physiology (28, 29). Two major proteins that function in cholesterol transport are NPC1 and NPC2. NPC2 is the central shuttle in a unidirectional transfer pathway that mobilizes cholesterol to NPC1, leading to NPC1 export of cholesterol from late endosomes (16, 23, 24, 30). Recently, it has been demonstrated that filoviruses, including Ebola and Marburg viruses, utilize NPC1 as intracellular receptors for entry (18). Interestingly, in addition to Ebola virus (family: *Filoviridae*), hepatitis C virus (family: *Flaviviridae*) entry also requires the cholesterol trafficking receptor Niemann-Pick C1-Like 1 (NPC1L1) (31), a NPC1 paralog which serves in cellular cholesterol absorption and homeostasis on the apical surface of intestinal enterocytes and human hepatocytes (20, 32). Moreover, the release of HIV-1 (family: *Retroviridae*) (33) and Chikungunya virus (family: *Togaviridae*) (27) is impaired in cells from patients with NPC disease. U18666A or imipramine, which mimics a NPC-deficient phenotype, also strongly inhibits the replication of several *flaviviridae* family members, including Zika, West Nile, and Dengue viruses (27). In our study, we present evidence that baculovirus utilizes the NPC1 homolog to enter into insect cells. Although the enveloped viruses described above belong to distinct viral families *(Baculoviridae*, *Filoviridae*, *Togaviridae* and *Flaviviridae)*, and possess different types (dsDNA, +ssRNA or -ssRNA) and sizes of viral genomes, they invariably hijack NPC1 in the cholesterol transport pathway to initiate infection. We speculate that NPC1 is the common entry receptor for many enveloped viruses during cellular infection. However, it should be noted that infection with some enveloped viruses from *Rhabdoviridae* (vesicular stomatitis virus) (18) or *Orthomyxoviridae* (infiuenza A virus) (34) occurs independently of NPC1.

However, despite both Ebola virus and baculovirus requiring NPC1 for viral entry, NPC1-dependent baculovirus infection of insect cells exhibits unique features. The EBOV glycoprotein GP is a Class I fusion protein. During virus entry, Ebola virus is internalized into host cells by receptor binding-mediated macropinocytosis and is then transported to late endosomes. In the endosome, cathepsin B-mediated cleavage removes mucin and glycan cap domains from the GP protein and exposes a hydrophobic cavity, which can mediate binding to the hNPC1 C domain. The specific binding between hNPC1 and GPcl was hypothesized to facilitate fusion between vial membrane and host endosomal membranes (23). In contrast, the BmNPV glycoprotein gp64 is a Class III fusion protein, and viral fusion is triggered by a caustic pH without proteolytic cleavage. We show here that BmNPC1 proteins are located at both the cell plasma membrane and intracellular compartments, and that BmNPC1 is capable of binding to gp64 at a neutral pH, indicating that BmNPC1 may serve as a host factor to facilitate virus attachment to the cell surface. The enhanced binding between BmNPC1 and gp64 at a lower pH as demonstrated in our study suggests that low pH triggered-conformational changes in the gp64 protein may expose certain epitopes or domains to allow better accessibility by BmNPC1, and the specific binding between gp64 and BmNPC1 may ultimately enable gp64 to maintain a certain structure which is essential for targeting its fusion domain into the endosomal membranes. However, the detailed mechanism of BmNPC1 protein in baculovirus entry will rely on future crystallographic examination of the BmNPC1 and gp64 complex. Such studies will ultimately shed light on how a wide range of enveloped viruses utilize the shared host factor NPC1 as part of fusion triggers for intracellular entry.

Baculovirus can transduce a broad range of cells including both vertebrate and invertebrate cells, suggesting that conserved host factors may be involved in viral entry. In our study, we have only examined the role of NPC1 in viral entry in insect cells; it will be of great interest in the future to assess whether NPC1 is involved in viral entry in mammalian cells. Furthermore, the cell surface localization of BmNPC1 in insect cells raises the possibility that BmNPC1 may act as a host factor for baculovirus in insect cells; future investigation to identify the role(s) of BmNPC1 in various steps of viral entry is warranted. In summary, our study demonstrated that BmNPV entry requires the cholesterol transporter BmNPC1 as the host factor for baculovirus infection, summarized with a hypothetical model (Fig. 6). In this model, baculovirus binds to the cell surface with the possibility of using BmNPC1 as one of the attachment factors, and is then internalized and trafficked to late endosomes. In the caustic pH environment of endosomes, gp64 undergoes conformational changes to expose certain domain(s) for better access by BmNPC1. The specific binding between gp64 and BmNPC1 subsequently facilitates viral fusion events to allow nucleocapsids to be released into the cytoplasm. Our work fills an important gap in baculovirus research, and identifies a new antiviral target against baculovirus infection. Elucidation of baculovirus entry mechanisms will further facilitate the application of baculovirus systems in eukaryotic gene delivery. As the cholesterol transporter NPC1 is shared by several viral families, this work provides a new avenue of inquiry that NPC1 may represent a common entry factor for many enveloped viruses entry.

**FIG 6.**
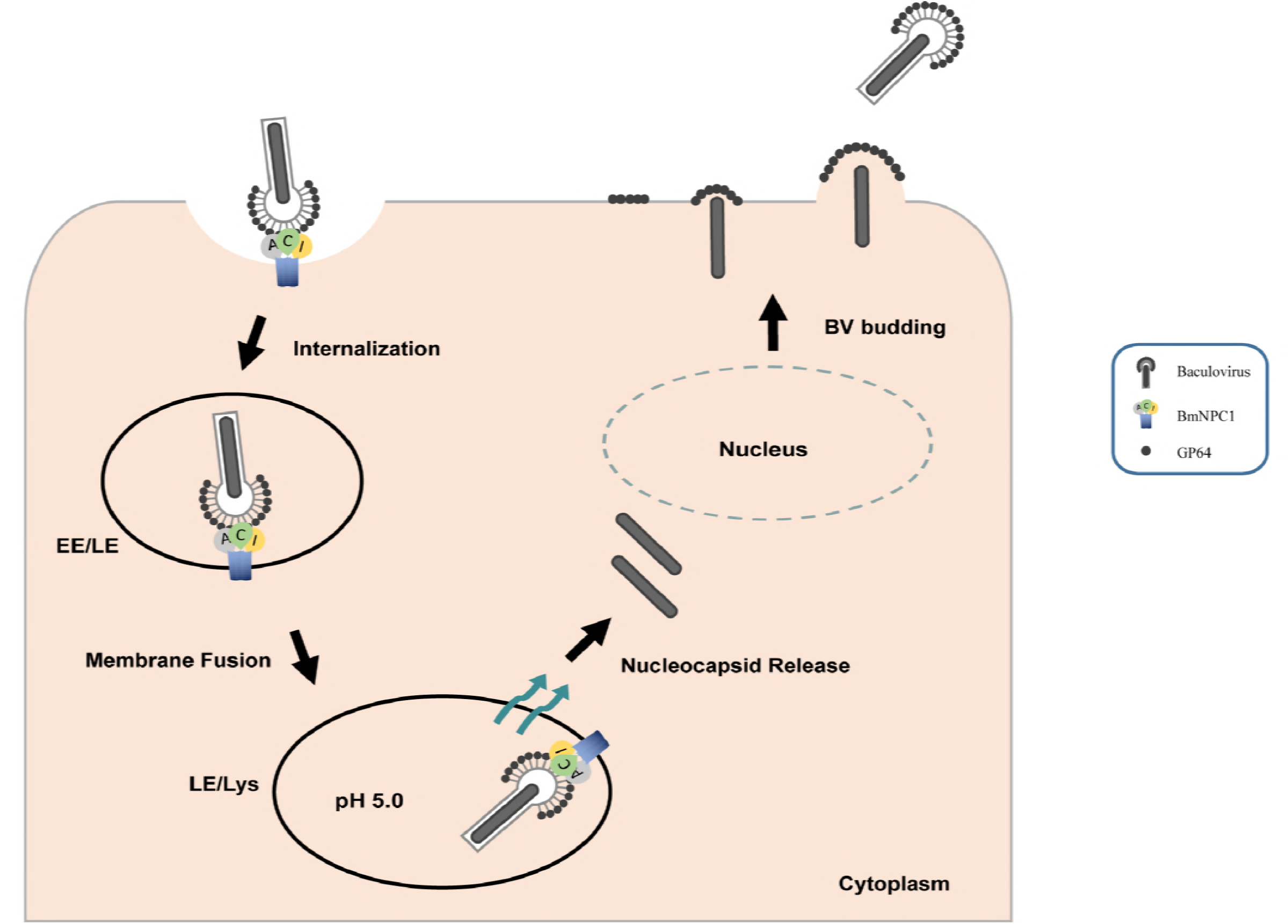
A model illustrating the role of BmNPC1 in baculovirus entry. Virus particles adhere to the plasma membrane by binding to NPC1, NPV gp64 could interact with BmNPC1-C domain, and trafficked to endosome (early/late endosomal, EE/LE) compartments, then the conformational change of gp64 were triggered by low pH, initiated the fusion of viral and host membranes and the cytoplasmic release of the viral nucleocapsid. After viral transcription and replication occur in the host cell nucleus, new BV particles are budded out from the basolateral membrane to spread the infection systemically.

## MATERIALS AND METHODS

### Cell culture and recombinant virus

*B. mori* cell line BmE cells were maintained at 28°C in Grace’s medium (Thermo Fisher Scientific, USA) supplemented with 10% (V/V) fetal bovine serum (FBS) (Thermo Fisher Scientific, USA) and 1%(V/V) penicillin-streptomycin (35). The recombinant BmNPV virus bearing an EGFP gene under the control of polyherin promoter was constructed by Bac-to-Bac Baculovirus Expression System according to the manufacturer’s protocol (Invitrogen, USA). The expression of EGFP can act as a reporter for monitoring viral gene expression and viral replication (36). Recombinant viruses were propagated in BmE cells and viral titers were measured by 50% tissue culture infectious dose (TCID_50_) based on EGFP expression as described (37).

### Cloning, protein expression, and purification of BmNPC1 and antibody production

The BmNPC1 transmembrane domain was predicted based on the website (http://www.cbs.dtu.dk/services/TMHMM-2.0/), and functional domains were predicted based on the website (http://smart.embl-heidelberg.de/). The full coding region of BmNPC1 was PCR amplified with specific primer (Table S1) from a BmE cell cDNA library and then cloned into the pMD19-T vector for sequencing. The sequences of BmNPC1 extracellular loop A (31 to 264 amino acids) or I (881 to 1152 amino acids) were amplified with primer pairs BmNPC1-A-PE and BmNPC1-I-PE (Table S1) and were then cloned into pET32a (+) plasmid for protein expression in *E. coli*. The recombinant BmNPC1-A, BmNPC1-I proteins with His tag were purified with a HisTrap HP 5 ml column (GE Healthcare, USA) according to the manufacturer’s instructions. The purified recombinant protein BmNPC1-A and BmNPC1-I were later used as an immunogen to immunize 7-week-old BALB/c mice with Freund’s complete adjuvant (Sigma-Aldrich, USA). The TDPVELWASPTSRS polypeptide from BmNPC1 (409 to 422 amino acids) coupled with the KLH tag were synthesized by GenScript and used to immunize New Zealand rabbits with Titermax Gold adjuvant. The anti-BmNPC1-A and anti-BmNPC1-I mouse serum, anti-BmNPC1 rabbit serum, and control sera from both naïve mice and rabbits immunized with KLH tag only were collected and stored at -20°C until use.

### BmNPC1-RNAi construction and transfection in BmE cells

The RNAi target prediction of BmNPC1 was performed according to the website (http://jura.wi.mit.edu/bioc/siRNA); two regions designated npc1-a and npc1-b were chosen based on scores for subsequent use. The sense and antisense fragments of npc1-a and npc1-b were amplified from the BmE genome with primer pairs npc1-RNAi-a and npc1-RNAi-b (Table S1), and were subsequently cloned into the RNAi vector pSL1180 with A3 intron as a spacer (3) (Fig 3A), generating RNAi vectors psl-BmNPC1-a and psl-BmNPC1-b. Unmodified pSL1180 vector designated as psl-null was used as a negative control for RNAi.

BmE cells (7 × 10^5^ cells/well) grown in six-well culture plates were transfected with 2 μg of psl-BmNPC1-a or psl-BmNPC1-b, or negative control psl-null siRNA, using 2 μl of X-treme GENE HP DNA tranfection reagent (Roche, Germany) following the manufacturer’s protocol for 48h before cells were subjected to viral infection. The knockdown effect of RNAi transfection was confirmed by mRNA expression via qPCR with primer set BmNPC1-q (Table S1).

### Construction and verification of NPC1-knockout (ΔNPC1) BmE cells

The CRISPR/Cas9 genomic editing tool was used to generate the stable BmNPC1-knockout BmE cell line. The Cas9 expression vector (named pSL1180-IE1-Cas9) and sgRNA expression vector (named PUC57-U6-gRNA) were constructed as previously described (38) (Fig S3A). The primer sequences generating the guide RNA (gRNA) targeting the BmNPC1 genes were designed based on the CRISPR website (http://crispr.dbcls.jp/); all candidate sgRNA target sequences bear the GN19NGG sequence. Next, extraction of two optimal target sites (gRNA-1 and gRNA-2) based on the CRISPR website (http://crispr.dbcls.jp/) was performed; the forward and reverse primers of the target sequence start with AAGT and AAAC, respectively. The corresponding oligos were annealed and ligated into a sgRNA expression vector which was digested by *Bbs* I to generate sgRNA-1 and sgRNA-2 expression plasmids. All sgRNA expression plasmids were sequenced by M13 primers for verification. For construction of the BmNPC1 replacement donor vector, homologous arms targeted upstream of the gRNA-1 site and downstream of the gRNA-2 site were amplified from *Bombyx mori* genomic DNA and designated as Left-HR and Right-HR, respectively. The selective markers *ie*1-DsRed-sv40 and A3-*neo*-sv40 were maintained in our laboratory. The Left-HR, DsRed expression cassettes, neomycin (neo) expression cassettes, and Right-HR were sequentially cloned into the pBluescript II KS (-) vector to generate the donor plasmid pKS-donor (Fig S3B). All clones were verified by sequencing of plasmids. The primers are listed in Supplemental Table S1.

### Reverse transcriptase PCR and Quantitative real-time PCR (qPCR)

Total RNA from BmE cell was extracted using Total RNA kit II (Omega, USA) following manufacturer’s protocol, and reverse transcription was carried out using MLV Reverse Transcriptase (Promega, USA). These cDNA samples were used to detect transcripts of BmNPC1 using the primers BmNPC1-q (Table S1). The BmRPL3 (gi|112982798) amplified with primers BmRPL3-q (Table S1) was used as the internal reference. Sample analysis was performed on the CFX96^TM^ Real-Time System (Biorad, France).

The viral DNA loads were calculated based on qPCR of the BmNPV gp41 gene. Total DNA from each sample was prepared with a Wizard Genomic DNA extraction Kit (Promega, USA) according to the manufacturer’s protocol, and the qPCR was performed using primers gp41-q (Table S1) targeting a 120-bp region of the BmNPV gp41 gene (AAC63752.1). The BmGAPDH gene amplified by primers BmGAPHG-q (Table S1) was used as the reference.

### Viral infection and quantification of infectivity

BmE cells or ΔNPC1-BmE cells (7 × 10^5^ cells/well) seeded into six-well culture plates were inoculated with BmNPV-GFP virus at an MOI of 1 for 2 h, after which cells were washed three times with serum-free Grace medium followed by incubation with Grace medium supplemented with 10% FBS. Viral infectivity was monitored by GFP expression positive cells at the indicated time points. The mean percentage of GFP positive cells from five representative fields was calculated, and the data presented are representative of three independent experiments. Next, the ΔNPC1-BmE cells were collected based on DsRed expression by flow cytometry (Fig S3C), and viral DNA loads were determined by relative quantification of viral gp41 gene using qPCR and were expressed as the fold of change compared to the corresponding control. Viral titers in RNAi-NPC1 cells were determined with a TCID_50_ endpoint dilution assay based on the GFP expression positive cells.

### U18666A and imipramine treatment

BmE cells (2 × 10^5^ cells/well) were seeded into twelve-well culture plates and pre-incubated with vehicle or various concentrations of imipramine dissolved in methanol (25, 50, 75 or 100 μM) or U18666A dissolved in serum-free Grace medium (0.5, 1, 5 or 10 μM) for 2 or 24 h, respectively. Drug pre-treated cells were subjected to BmNPV-GFP virus infection at an MOI of 1 for 72 h in the presence of drug, and viral infectivity was evaluated by GFP positive cells and quantification of the relative amount of viral DNA in supernatants by qPCR.

### In vitro expression of BmNPC1 and gp64 proteins and Co-immunoprecipitation

Fragments of BmNPC1-A, BmNPC1-C, BmNPC1-I and BmNPV gp64 were amplified and cloned into pT7CFE1-NHis-GST-CHA vector (Thermo Fisher Scientific, USA) for subsequent expression by 1-Step CHO High-Yield In Vitro Translation (IVT) Kit following the manufacturer’s protocol (Thermo Fisher Scientific, USA). A, C, and I-domains of BmNPC1 were expressed with His_9_-tag and GST-tag to the N-terminus and a HA-tag to the C-terminus; gp64 protein were only expressed with His_9_-tag and GST-tag to the N-terminus (not include a HA-tag). The expression of recombinant proteins was confirmed by Western blot using anti-HA monoclonal antibody (Sigma-Aldrich, USA) or anti-gp64 monoclonal antibody (Abcom, UK). Co-immunoprecipitation was performed in accordance with standard protocols to probe the interaction between gp64 and BmNPC1 proteins. In brief, the protein G agarose beads (BioRad, USA) bearing gp64 protein through mouse anti-gp64 antibody were incubated with NPC1-A, NPC1-C or NPC1-I protein overnight at 4°C with gentle shaking. Proteins eluted from the beads were probed with anti-HA monoclonal antibody via Western blot. Co-immunoprecipitation was also performed in a reciprocal manner; in which NPC1-A, NPC1-C, and NPC1-I conjugated beads were incubated with gp64 protein, with proteins eluted from the beads probed with mouse anti-gp64 monoclonal antibody.

### Yeast two-hybrid assay

The yeast two-hybrid assay was used to confirm the interaction between NPC1-A, NPC1-C, NPC1-I and gp64 in vivo according to the previously described (39). The bait and prey constructs pairs pGBKT7-gp64/pGADT7-NPC1-A, pGBKT7-gp64/pGADT7-NPC1-C, pGBKT7-gp64/pGADT7-NPC1-I, and pGBKT7-NPC1-C /pGADT7-gp64 were transformed simultaneously into competent yeast cells to examine the protein interaction. All primers used are listed in Supplemental Table S1.

### Immunofluorescence assay (IFA)

BmE cells grown on cover glasses in 12-well plates (Corning, USA) for 12h at 28°C, then were fixed by 4% paraformaldehyde at room temperature for 30 min followed by permeabilization with 0.2% Triton X-100 or without permeabiliztion. The cells were stained with PcAb-hNPC1 antibody (Abcom, UK) or negative rabbit IgG for immunofluorescence microscopy as described previously (40), and imaged with an Olympus confocal microscope.

### Antibody blocking assay

BmE cells (2 × 10^5^ cells/well) seeded in twelve-well culture plates were incubated with one of the following antibodies: mouse anti-NPC1-A,anti-NPC1-I polyclonal antibody, rabbit anti-NPC1-C polyclonal antibody, mouse or rabbit naïve serum, at a concentration of 10 μg/ml for 2h at 28°C, after which cells were infected with BmNPV-GFP at an MOI of 3. At 72 h p. i. infected cells were fixed for microscopy and the supernatants were harvested for measuring viral DNA load by qPCR as described previously.

### Statistics

Independent-samples T tests were used for statistical analysis. Significant differences are marked with * at *P* < 0.05, ** at *P* < 0.01 and *** at *P* < 0.001; n.s, non-significant, respectively. All results are graphed as means ± SD for triplicate samples. All the data presented are representative of a minimum of three independent experiments.

### Ethics statement

All animal experiments were conducted in accordance with Laboratory Animals Ethics Review Committee of Southwest University guidelines (Chongqing, China) with committee approval for this study (Permit Number: AERCSWU2017-7). Mice were maintained in accordance with recommendations of the committee for the purpose of control and supervision of experiments on animals.

## ACKNOWLEDGMENTS

This work was supported by the National Natural Science Foundation of China (grant number 31770160, 31470250, 31270138, 31702185) and Fundamental Research Funds for the Central Universities (XDJK2018AA001, XDJK2015A010).

